# Impaired Therapeutic Efficacy of Bone Marrow Cells from Post-Myocardial Infarction Patients in the TIME and LateTIME Clinical Trials

**DOI:** 10.1101/721860

**Authors:** Xiaoyin Wang, Ronak Derakhshandeh, Hilda J. Rodriguez, Daniel D. Han, Dmitry S. Kostyushev, Timothy D. Henry, Jay H. Traverse, Lem Moyé, Robert D. Simari, Doris A. Taylor, Matthew L. Springer

## Abstract

Implantation of bone marrow-derived cells (BMCs) into mouse hearts post-myocardial infarction (MI) limits cardiac functional decline. However, clinical trials of post-MI BMC therapy have yielded conflicting results. While most laboratory experiments use healthy BMC donor mice, clinical trials use post-MI autologous BMCs. Post-MI mouse BMCs are therapeutically impaired, due to inflammatory changes in BMC composition. Thus, therapeutic efficacy of the BMCs progressively worsens after MI but recovers as donor inflammatory response resolves. The availability of post-MI patient BM mononuclear cells (MNCs) from the TIME and LateTIME clinical trials enabled us to test if human post-MI MNCs undergo a similar period of impaired efficacy. We hypothesized that MNCs from TIME trial patients would be less therapeutic than healthy human donor MNCs when implanted into post-MI mouse hearts, and that therapeutic properties would be restored in MNCs from LateTIME trial patients. Post-MI SCID mice received MNCs from healthy donors, TIME patients, or LateTIME patients. Cardiac function improved considerably in the healthy donor group, but neither the TIME nor LateTIME group showed therapeutic effect. Conclusion: post-MI human MNCs lack therapeutic benefits possessed by healthy MNCs, which may partially explain why BMC clinical trials have been less successful than mouse studies.

## Introduction

Autologous cell therapy after myocardial infarction (MI), for the purpose of repairing damaged regions or preserving tissue at risk, is the focus of intense research and debate. Delivery of various populations of bone marrow-derived cells (BMCs) to the infarcted myocardium has been reported to improve post-MI cardiac function in most preclinical experiments, despite debate about the underlying mechanisms^1^. We and others have shown that secreted or intracellular products from various populations of BMCs mediate robust therapeutic benefits to post-MI cardiac function in rodent models^2-6^. However, positive preclinical outcomes with autologous cells across species have not translated into approved human cell therapies, as the resulting cardiac functional improvement in humans after MI are modest at best^7-10^.

An important difference between rodent and human situations is that human patients undergoing autologous BMC therapy are middle aged or older and have multiple risk factors including a recent MI. In contrast, rodent BMC therapy cannot be autologous and involves distinct donors that are typically healthy and young. We previously reported that both the post-MI state and advanced age impair the therapeutic properties of BMCs in rodents, in each case involving reduction in the number of B-lymphocytes in bone marrow^2,11,12^. Notably, the impairment of therapeutic efficacy in post-MI BMCs involves a limited duration inflammatory response, the inhibition of which prevents the therapeutically impaired state in donor BMCs^12^. We further showed that the timing of the appearance and resolution of the post-MI acute inflammatory response correlated with a decline and recovery in BMC therapeutic efficacy, with a progressive impairment of BMCs observed over the first week post-MI, followed by a partial return to normalcy by 21 days post-MI. However, in humans, this has not been evaluated.

In the TIME (NCT00684021) and LateTIME (NCT00684060) clinical trials conducted by the NHLBI Cardiovascular Cell Therapy Research Network (CCTRN), patients with ST-elevation myocardial infarction (STEMI) were treated with autologous bone marrow mononuclear cells (BM MNCs) harvested 3 and 7 days post-MI (TIME trial)^10^ or 2-3 weeks post-MI (LateTIME trial)^9^. However, no therapeutic improvement in left ventricular ejection fraction (LVEF) was detected in either trial. To explore whether the therapeutic insufficiency of the MNCs in the clinical trials would also be evident in the mouse model, we implanted the same post-MI patient MNCs into post-MI immunodeficient mouse hearts, using the conditions of our previous experiments. This enabled us to test if the human post-MI MNCs undergo a similar period of impaired efficacy as we previously observed in post-MI mouse BMCs^12^ and if the therapeutic properties would be restored with time after MI. Here, we show that post-MI MNCs from these clinical trial patients, when harvested up to 3 weeks after MI, lack the therapeutic effects exhibited by healthy human BM MNCs when implanted into post-MI mouse hearts.

## Results

We hypothesized that human BM MNCs harvested at 3 or 7 days post-MI are therapeutically impaired and that human BM MNCs harvested 3 weeks post-MI are less impaired, based on results from our mouse-to-mouse experiments in which a progressive impairment of BMCs occurred over the first week post-MI, followed by a partial return to normalcy at 21 days post-MI^12^. To test this hypothesis, we surgically induced MI in SCID mice and injected them intramyocardially at 3 days post-recipient MI with cells from the following experimental conditions: pooled BM MNCs harvested 3 and 7 days post-MI from TIME patients (n=6, all male, mean age 46.0±4.7 (SD) years), pooled BM MNCs harvested 2-3 weeks post-MI from LateTIME patients (n=6, all male, 47.4±2.2 years), or commercially obtained but comparably prepared (see Methods) pooled age-matched healthy donor BM MNCs (n=6, 5 male + 1 female, 43.3±6.4 years) (see Table 1). A fourth group received vehicle negative control injections (HBSS; see Methods). Recipient mouse cardiac function represented by LVEF and end-systolic and diastolic volumes (ESV, EDV) was assessed by echocardiography. As expected, LVEF declined significantly over 28 days post-MI in the HBSS vehicle group and improved considerably in the healthy donor group. However, no improvement of cardiac function in mice was observed in the TIME or LateTIME groups (Fig. 1 and Table 2). Although the TIME group exhibited slightly better function than the vehicle group, the difference was not significant; and function in the LateTIME group was almost identical to that in the TIME group, suggesting that post-MI human BM MNCs harvested even after 3 weeks post-MI lack the therapeutic benefit that healthy human BM MNCs bestow on post-MI mice. Infarct size at 28 days post-MI trended toward larger values in the TIME, LateTIME, and HBSS vehicle groups compared to the healthy donor group, although variability was high and the differences did not reach significance.

**Table 1.**
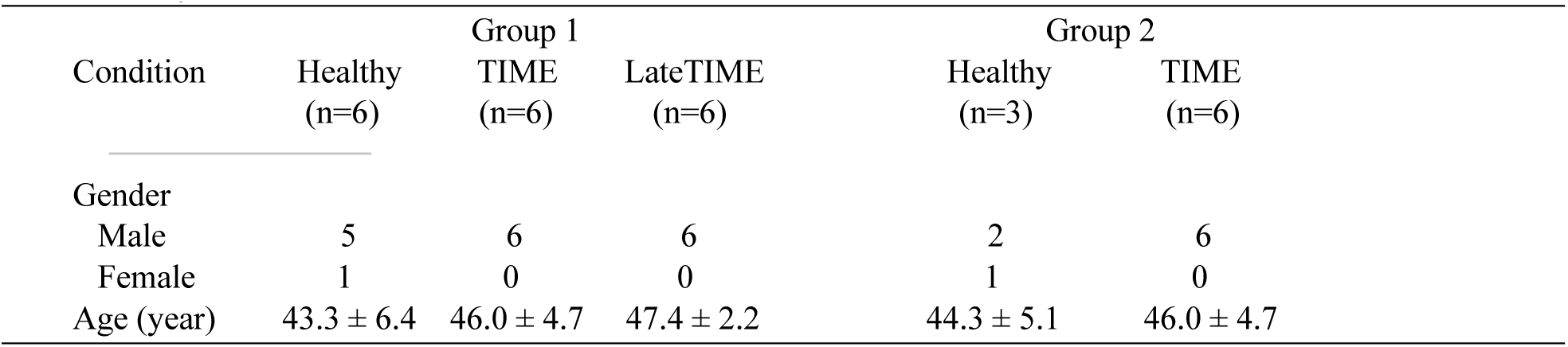
Subject information for bone marrow mononuclear cells.

**Table 2.**
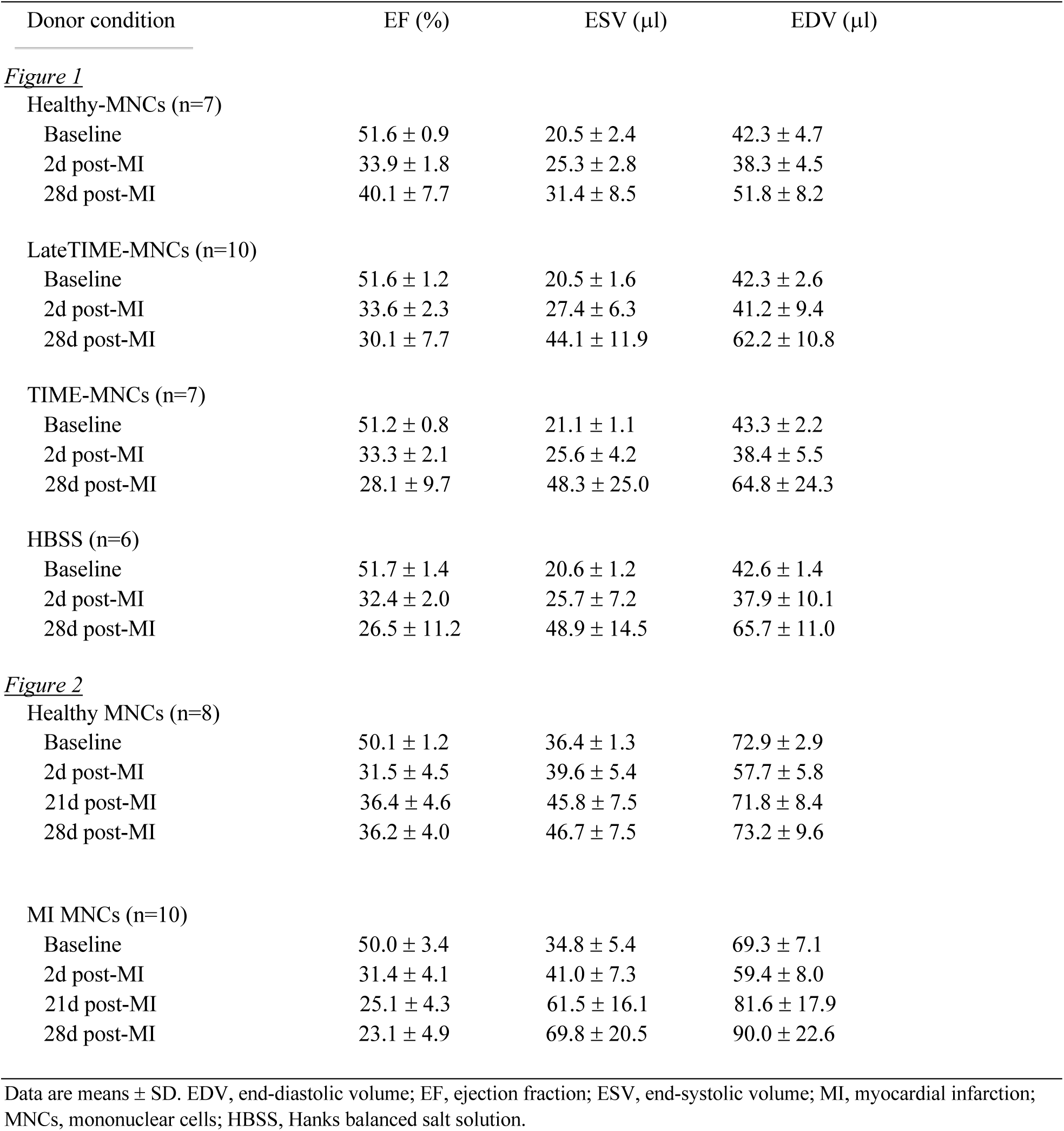
Recipient echocardiographic parameters pre-and post-MI.

| Donor condition | EF (%) | ESV (μl) | EDV (μl) |
| --- | --- | --- | --- |
| *Figure 1* |  |  |  |
| Healthy-MNCs (n=7) |  |  |  |
| Baseline | 51.6 ± 0.9 | 20.5 ± 2.4 | 42.3 ± 4.7 |
| 2d post-MI | 33.9 ± 1.8 | 25.3 ± 2.8 | 38.3 ± 4.5 |
| 28d post-MI | 40.1 ± 7.7 | 31.4 ± 8.5 | 51.8 ± 8.2 |
| LateTIME-MNCs (n=10) |  |  |  |
| Baseline | 51.6 ± 1.2 | 20.5 ± 1.6 | 42.3 ± 2.6 |
| 2d post-MI | 33.6 ± 2.3 | 27.4 ± 6.3 | 41.2 ± 9.4 |
| 28d post-MI | 30.1 ± 7.7 | 44.1 ± 11.9 | 62.2 ± 10.8 |
| TIME-MNCs (n=7) |  |  |  |
| Baseline | 51.2 ± 0.8 | 21.1 ± 1.1 | 43.3 ± 2.2 |
| 2d post-MI | 33.3 ± 2.1 | 25.6 ± 4.2 | 38.4 ± 5.5 |
| 28d post-MI | 28.1 ± 9.7 | 48.3 ± 25.0 | 64.8 ± 24.3 |
| HBSS (n=6) |  |  |  |
| Baseline | 51.7 ± 1.4 | 20.6 ± 1.2 | 42.6 ± 1.4 |
| 2d post-MI | 32.4 ± 2.0 | 25.7 ± 7.2 | 37.9 ± 10.1 |
| 28d post-MI | 26.5 ± 11.2 | 48.9 ± 14.5 | 65.7 ± 11.0 |
| *Figure 2* |  |  |  |
| Healthy MNCs (n=8) |  |  |  |
| Baseline | 50.1 ± 1.2 | 36.4 ± 1.3 | 72.9 ± 2.9 |
| 2d post-MI | 31.5 ± 4.5 | 39.6 ± 5.4 | 57.7 ± 5.8 |
| 21d post-MI | 36.4 ± 4.6 | 45.8 ± 7.5 | 71.8 ± 8.4 |
| 28d post-MI | 36.2 ± 4.0 | 46.7 ± 7.5 | 73.2 ± 9.6 |
| MI MNCs (n=10) |  |  |  |
| Baseline | 50.0 ± 3.4 | 34.8 ± 5.4 | 69.3 ± 7.1 |
| 2d post-MI | 31.4 ± 4.1 | 41.0 ± 7.3 | 59.4 ± 8.0 |
| 21d post-MI | 25.1 ± 4.3 | 61.5 ± 16.1 | 81.6 ± 17.9 |
| 28d post-MI | 23.1 ± 4.9 | 69.8 ± 20.5 | 90.0 ± 22.6 |

Data are means ± SD. EDV, end-diastolic volume; EF, ejection fraction; ESV, end-systolic volume; MI, myocardial infarction; MNCs, mononuclear cells; HBSS, Hanks balanced salt solution.

**Figure 1.**
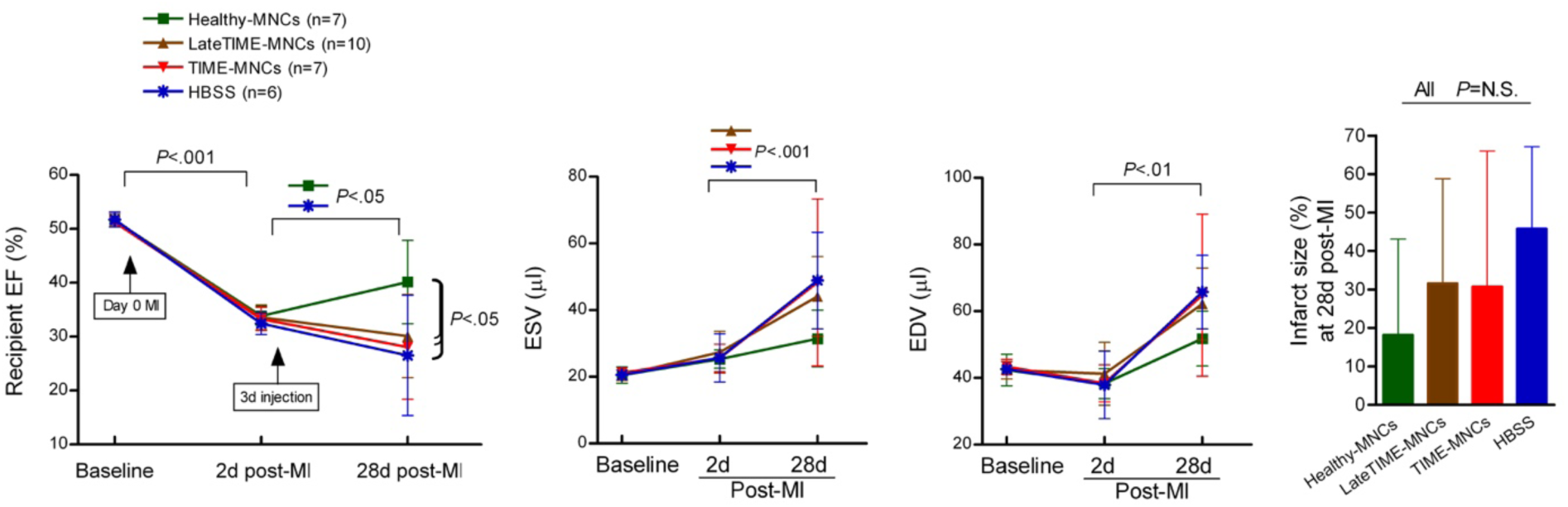
Impaired therapeutic efficacy of post-MI patient BM MNCs from the TIME and LateTIME clinical trials. Post-MI MNCs were similar to HBSS vehicle control in their lack of beneficial effect on recipient cardiac function, whereas healthy MNCs improved LVEF and minimized ventricular chamber dimensions such that ESV in only the healthy donor group was not significantly increased, with a non-significant difference between EDV from recipients of healthy cells vs. the other groups. Infarct size at 28 days post-MI showed a trend toward higher values in the post-MI donor groups and HBSS vehicle group as compared to the healthy donor group, although no significant differences were reached (*P* = 0.925 – 1.00). Error bars = SD.

While this experiment was performed blinded with direct comparison of the four implantation conditions, a previous, non-blinded pilot experiment yielded similar results and is provided here to demonstrate reproducibility. A pool of MNCs from the same 6 TIME patients as in the Figure 1 experiment was similarly implanted into one group, compared to a pool of MNCs from 3 healthy donors implanted into another group (Fig. 2 and Table 2). In this case, LV function was measured at both 21 and 28 days post-MI. The difference in function between the two groups was comparable to that observed in the Fig. 1 experiment, and we additionally observed that ESV and EDV were stable from day 21 to day 28 in the healthy donor MNC group, but continued to worsen during that time in the post-MI TIME MNC group.

**Figure 2.**
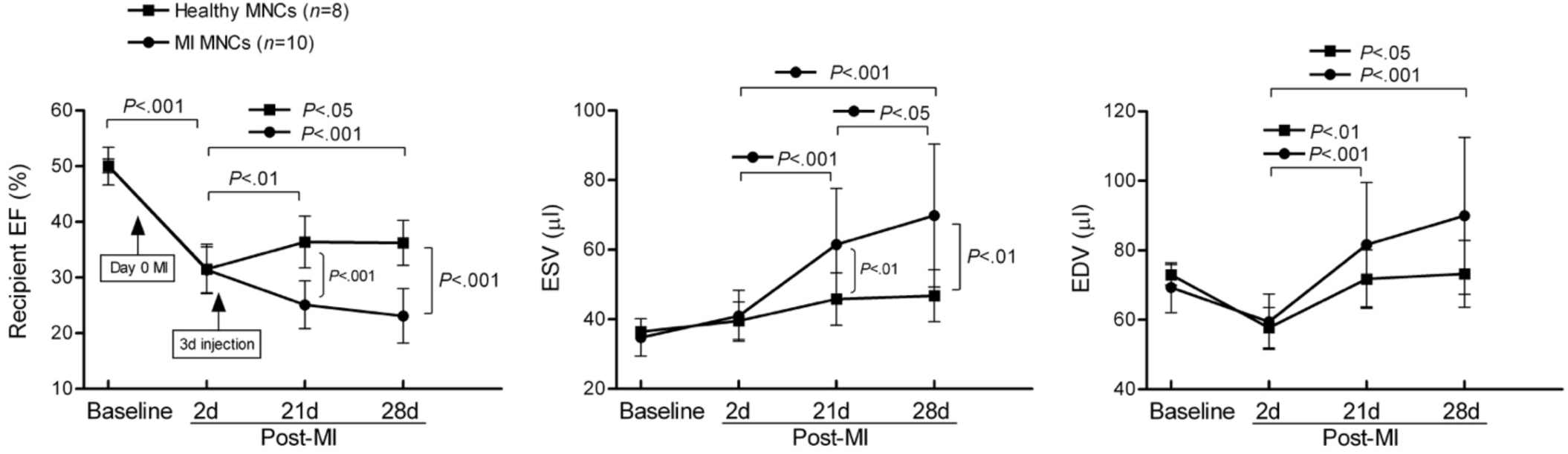
Reproducibility of TIME MNC therapeutic impairment and comparison of day 21 and 28 function. Therapeutic impairment of the TIME MNCs was also apparent in this earlier (non-blinded) pilot experiment. Moreover, EF was stable between days 21 and 28 post-recipient MI in the healthy donor MNC group, but declined during that period in the TIME MNC group. Error bars = SD.

Therefore, both experiments were consistent in demonstrating that BM MNCs from middle-aged post-MI patients were therapeutically impaired relative to age-matched healthy donor MNCs when implanted into hearts of 3 day post-MI SCID mice.

## Discussion

Our previous preclinical findings from implantation of post-MI mouse BMCs^12^ suggested that human post-MI BM MNCs from the TIME trial, and possibly from the LateTIME trial, would be therapeutically impaired when implanted into post-MI immunodeficient mouse hearts. Consistent with this hypothesis, autologous MNCs obtained from TIME and LateTIME patients were not beneficial in the respective clinical trials^9,10^, and our present study indicates that they lack the therapeutic properties of healthy donor MNCs when implanted into mouse hearts.

We reported previously that at 3 days post-MI, the recipient mouse heart was responsive to BMC therapy, but that BMC implantation at 7 and 14 days post-MI did not result in optimal functional improvement^13^, potentially because the local inflammatory state of the myocardium somehow interfered with the beneficial effects caused by implantation of intact or lysed cells. However, we subsequently identified 3 days post-MI in the donor as the time at which the BMCs are the least effective^12^, with 21 days shown to be a more ideal donor time frame – presenting a conundrum for effective autologous cell therapy in which the most effective conditions would involve ∼21 day post-MI BMCs implanted into ∼3 day post-MI hearts. We proposed that the same temporal mismatch may underlie negative results of autologous human trials such as TIME and LateTIME. Even trials such as REPAIR-AMI, in which therapeutic effects have been reported^7^, may have been hindered by this effect.

Notably, the gradual decline and recovery of therapeutic efficacy of the BMCs after MI correlated in time with the appearance and resolution of the post-MI acute inflammatory response^12^. Consequently, based on the results from our mouse-to-mouse experiments, a possible explanation for the negative results from post-MI patient BMCs to MI mice is that the 3-7 day post-MI BMCs in the TIME trial were still in the inflammatory state and problematic for therapy. While the recovery by day 21 post-MI led us to hypothesize that the 2-3 week post-MI BMCs in the LateTIME trial would give better results than TIME trial, that effect was not evident here. Interestingly, the LateTIME group exhibited slightly better values for EF, ESV, and EDV than the TIME and HBSS groups, but the differences were minor and non-significant. The hypothetical increase in BMC functional ability after 3 weeks post-MI may have been simply inadequate to cause detectable therapeutic benefit. It is quite possible that the difference in timing of recovery in mice vs. humans^14^ makes 3 weeks post-MI in the LateTIME patients still too early to observe the recovery of therapeutic properties. Additionally, our mouse results are based on a permanent ligation MI model rather than the ischemia/reperfusion model that more closely mimics primary percutaneous coronary intervention treatment of STEMI as utilized in almost all clinical trials using BMCs. Regardless, our findings underscore the importance of understanding MI-induced BMC impairment and developing ways to limit or prevent it if autologous BMC therapy is to be pursued as a viable clinical approach.

Regarding a mechanistic explanation of why BMCs are therapeutically impaired after MI, findings from our group^12^ and others^15,16^ suggest that acute inflammation resulting from the MI alters whole BMC composition, particularly reducing the level of bone marrow B lymphocytes, which appear to mediate therapeutic activity in this and other models of cardiac and cerebral injury^2,12,17-20^. We recently reported that partial depletion of B cells from young healthy donor mouse BMCs reduces the BMCs’ therapeutic efficacy, and that injection of healthy intact B cells or B cell lysate into post-MI myocardium can bestow similar therapeutic effects as whole BMCs or BMC lysate^2^. This suggests that the reduction of the number of B cells in the bone marrow is a consequence of both age and/or MI that limits therapeutic efficacy of autologous BMC therapy. Supporting this notion, follow-up studies of cell product and of peripheral blood from the CCTRN trials have revealed that the level of bone marrow B cells from the individual patients from TIME trial, as well as the FOCUS heart failure trial, correlated positively with their individual clinical outcomes^21,22^.

Our study has several limitations. First, we experienced a surprisingly high spontaneous mortality rate in these SCID mice that we have not experienced in previous SCID mouse experiments, which severely reduced our group size. We were unable to simply replace animals and inject more cells because they were pooled from unique single patient-specific aliquots and were not replaceable (the n values in the graphs refer to recipient mouse group size). However, despite the smaller-than-expected group size, our results fortunately were significant.

Second, our conclusion from these experiments is that BM MNCs from TIME and LateTIME trials lacked therapeutic properties possessed by BM MNCs from healthy donors. However, we acknowledge that the preparations of cells were not 100% identical. BM MNCs from the TIME and LateTIME trials were isolated at each CCTRN clinical site by an automatic Ficoll separation protocol (Sepax), whereas the healthy BM MNCs were isolated by a manual Ficoll protocol at AllCells. Nonetheless, cell viability was good in all samples. Still, Assmus and colleagues reported that contamination of BM MNCs with red blood cells (RBCs) correlated with reduced recovery of LVEF in patients treated with the MNCs^23^. Because Sepax separation can yield BM MNCs with a greater degree of RBC contamination than manual Ficoll separation techniques, the RBC contamination could explain at least part of the reduced therapeutic effects of the Sepax-separated TIME and LateTIME trial BM MNCs relative to the manually isolated healthy donor BM MNCs. However, in all of our samples before implantation, RBCs were lysed and the remaining cells were rinsed, so it is unlikely that RBCs were co-implanted with the clinical trial samples. Furthermore, while the RBCs may have caused lasting damage to the MNCs before the RBCs were lysed, the MNCs from all donors (healthy and patient) would have been exposed to RBCs in the bone marrow and/or during the BM harvest before density gradient enrichment. Therefore, the brief disparate durations of co-existence between RBCs and MNCs during cell enrichment before RBC lysis and rinsing would be unlikely to explain the profound difference in therapeutic efficacy between the patient and healthy donor MNCs.

Lastly, while we did the best we could to gender/age match the healthy subjects to heart disease patients, the patients were all male but our healthy group included one female (the abundance of male subjects was an unavoidable result of our attempt to age-match the groups and to avoid elderly patients, as we have also shown advance age to impair BMC therapeutic efficacy^2,11^).

In conclusion, post-MI human BM MNCs lack the therapeutic benefit that healthy human MNCs bestow on post-MI mice, when harvested up to 3 weeks after MI. This may partially explain why BMC therapy clinical trials have been less successful than rodent studies.

## Methods

### Animals

All animal procedures were approved by the Institutional Animal Care and Use Committee of the University of California, San Francisco and performed in accordance with the recommendations of the American Association Accreditation of Laboratory Animal Care. Male SCID mice (CB17SC-sp/sp; 8 weeks) were obtained from Taconic Biosciences (Germantown, NY) and subjected to experiments as recipients at 9-10 weeks. Target group size was 10 if enough donor cells were available.

### Surgical induction of myocardial infarction

MI was surgically induced through permanent coronary artery ligation as we have described previously^12^. Briefly, mice were subjected to the MI surgery under anesthetization with 2% isoflurane and received subcutaneous buprenorphine (0.1 mg/kg) for analgesia at the time of surgery and at the end of the day. The heart was exposed via a parasternotomy and the left anterior descending coronary artery was permanently ligated ∼2 mm below the tip of the left atrium. The total percentage of mortality after MI surgery was 25%.

### Preparation and implantation of human BM MNCs

Stored BM MNCs obtained from TIME and LateTIME clinical trial patients^9,10^ were utilized in these studies. These patients provided written informed consent for participation in the trials and for the use of their cells in future research. All methods pertaining to human subjects research were carried out in accordance with relevant guidelines and regulations, and both trials were approved by the following independent institutional review boards (IRBs) at each clinical center and the data coordinating center: Cleveland Clinic IRB, University Hospitals Case Medical Center IRB, St. Luke’s Episcopal Hospital IRB, IRB of Human Subjects Research for Baylor College of Medicine and Affiliated Hospitals, University of Florida IRB, Pepin Heart Hospital RERB, Abbott Northwestern Hospital IRB, University of Minnesota IRB, Mayo Clinic IRB, Vanderbilt University IRB, University of Texas-Houston IRB. These MNCs had been prepared from BM aspirate harvested at 3-5 or 7-9 days (TIME) or 14-21 days (LateTIME) post-MI, and isolated on Ficoll gradients using an automated closed system (Sepax)^24^. Healthy de-identified donor BM MNCs were purchased from AllCells (Alameda, CA). These MNCs were manually isolated at AllCells using Ficoll conditions like that in the Sepax technique used to isolate the TIME and LateTIME cells. All cell samples were cryopreserved and stored frozen in liquid nitrogen vapor phase. On the day of implantation, all aliquots to be pooled were thawed, washed, and resuspended in Hanks balanced salt solution (HBSS)/0.5% BSA for injection. Briefly, for each experimental group, one vial per patient or healthy donor was quickly thawed in a 37°C water bath, gently rinsed with prewarmed (37°C) medium (RPMI-1640 containing 10% FBS), pooled together, and centrifuged at 200 g for 15 min at room temperature. The supernatant was removed carefully and then the cell pellet was suspended in 15-20 ml of pre-warmed RPMI-1640/10% FBS and centrifuged again. After the two washes, the supernatant was removed and the pellet was resuspended in 5 ml of 1×RBC lysis buffer, mixed well, and incubated for 5 min at room temperature. After RBC lysis, the cell suspension was diluted with 20 ml of PBS/0.5% BSA/EDTA and centrifuged immediately at 500 g for 5 min at room temperature. The supernatant was aspirated and the pellet was resuspended with pre-warmed (37°C) buffer consisting of HBSS and 0.5% BSA for implantation. The cell concentration was adjusted to 10^8^ viable cells/ml. For MNC implantation into myocardium,10^6^ cells in HBSS/0.5% BSA were split into two 5 μl injections and implanted into myocardium by closed-chest ultrasound-guided injection using a Vevo660 micro-ultrasound system (VisualSonics Inc., Toronto) as we have described previously^25^, to target the infarct border zone and ensure successful implantation in the myocardial wall rather than the ventricular cavity. Each pool of BM MNCs was randomly allocated for 10 MI recipient mice per group, and investigators were blinded to the identity of MNCs during cell injection. Recipient mice were always injected at 3 days post-MI. Injection of HBSS/0.5% BSA served as a negative control.

### Measurements of cardiac function and infarct scar size

Serial echocardiography was performed in mice anesthetized with ∼1.25% isoflurane at baseline, 2 days post-MI (prior to injection), 21 days post-MI (for Figure 2 experiment only), and 28 days post-MI with the Vevo660 micro-ultrasound system. In the earlier pilot experiment, to compare changes in cardiac function post-MI, echocardiography was done at day 21 and day 28 post-MI. The ESV and EDV were obtained in two-dimensional mode at parasternal long axis view on echocardiography, and the LVEF was measured as we have previously described^26^. Echocardiography was performed and interpreted by a blinded investigator. Infarct scar size was calculated by a blinded investigator through histological measurement of the fibrotic scar zone using a midline arc length approach according to our published protocol^27^.

### Statistics

Power calculation based on standard deviations from within-group comparisons in our previous MI experiments determined that n=10/group was sufficient to detect changes in cardiac function at a power of 0.8 and significance level of 0.05. For comparisons involving multiple groups and times, we fit a 2-factor (treatment condition and time) repeated-measures analysis of variance to all the data at once using a mixed model estimated with restricted maximum likelihood estimation with an unstructured covariance matrix of residuals, then tested for differences over time and across treatment condition using contrasts and pairwise comparisons, adjusted for multiple comparisons using the Sidak method. Calculations were done with Stata 13.1.

## Data Availability

The datasets resulting from this study are available from the corresponding author upon reasonable request.

## Acknowledgements

We thank the NHLBI Cardiovascular Cell Therapy Research Network (CCTRN) for their support in this study and Judy Bettencourt, MPH of UTHealth for editorial assistance. This work was supported by NIH grants R01 HL086917, R21 HL097129, NIH cooperative agreement UM1 HL087318, and by American Heart Association grant 15GRNT225900001.

## Author Contributions

X.W. conducted experiments, helped conceive of and performed experiments and data analysis with participation of R.D., H.J.R., D.D.H., and D.K. X.W. and M.L.S. conceived of the project; T.D.H., J.H.T., L.M., R.D.S, and D.A.T. provided collaborative experimental support for the project. X.W. and M.L.S. wrote the manuscript, and D.A.T. provided critical revisions.

